# Phasic activation of dorsal raphe serotonergic neurons increases pupil-linked arousal

**DOI:** 10.1101/2020.06.25.171637

**Authors:** Fanny Cazettes, Davide Reato, João P. Morais, Alfonso Renart, Zachary F. Mainen

## Abstract

Variations in pupil size under constant luminance are closely coupled to changes in arousal state [1–5]. It is assumed that such fluctuations are primarily controlled by the noradrenergic system [6–9]. Phasic activity of noradrenergic axons precedes pupil dilations associated with rapid changes in arousal [7,9], and is believed to be driven by unexpected uncertainty [1,10–16]. However, the role of other modulatory pathways in the control of pupil-linked arousal has not been as thoroughly investigated, but evidence suggests that noradrenaline may not be alone [7,17,18]. Administration of serotonergic drugs seems to affect pupil size [19–23], but these effects have not been investigated in detail. Here, we show that transient serotonin (5-HT) activation, like noradrenaline, causes pupil-size changes. We used phasic optogenetic activation of 5-HT neurons in the dorsal raphe nucleus (DRN) in head-fixed mice locomoting in a foraging task. 5-HT-driven modulations of pupil size were maintained throughout the photostimulation period and sustained for several seconds after the end of the stimulation. The activation of 5-HT neurons increased pupil size additively with locomotor speed, suggesting that 5-HT transients affect pupil-linked arousal independently from locomotor states. We found that the effect of 5-HT on pupil size depended on the level of environmental uncertainty, consistent with the idea that 5-HT may report a salience or surprise signal [24]. Together, these results challenge the classic view of the neuromodulatory control of pupil-linked arousal, revealing a tight relationship between the activation of 5-HT neurons and changes in pupil size.

## RESULTS AND DISCUSSION

### Tracking pupil-linked arousal during a foraging task for head-fixed mice

We tracked pupillary fluctuations while head-fixed mice foraged for water. Mice had the choice to exploit either one of two resource sites (Figure 1A). Reward delivery at a given site was probabilistic (given by P_REW_) and switched stochastically to 0 after a variable number of licks, controlled by the probability of site depletion (P_DPL_, see STAR Methods and [25]). Mice were trained to remain still while licking at a given site and to run a set distance on a treadmill to switch between sites. Thus, the behavior consisted of periods of locomotion and stillness (including licking) of various lengths and onset timings (Figure 1B). Consistent with previous reports on pupil-linked arousal [2,26,27], we observed a tight relationship between pupil size and locomotor states (example in Figure 1C; across sessions, r = 0.42 ± 0.14, p<10^−7^, Figure S1). Specifically, transition from stillness to locomotion was accompanied by an increase in pupil size (Figure 1D). Pupil size also spontaneously varied during stillness epochs (Figure 1C, event 3, Figure S1), indicating periods of elevated arousal in absence of locomotion [27].

**Figure 1.**
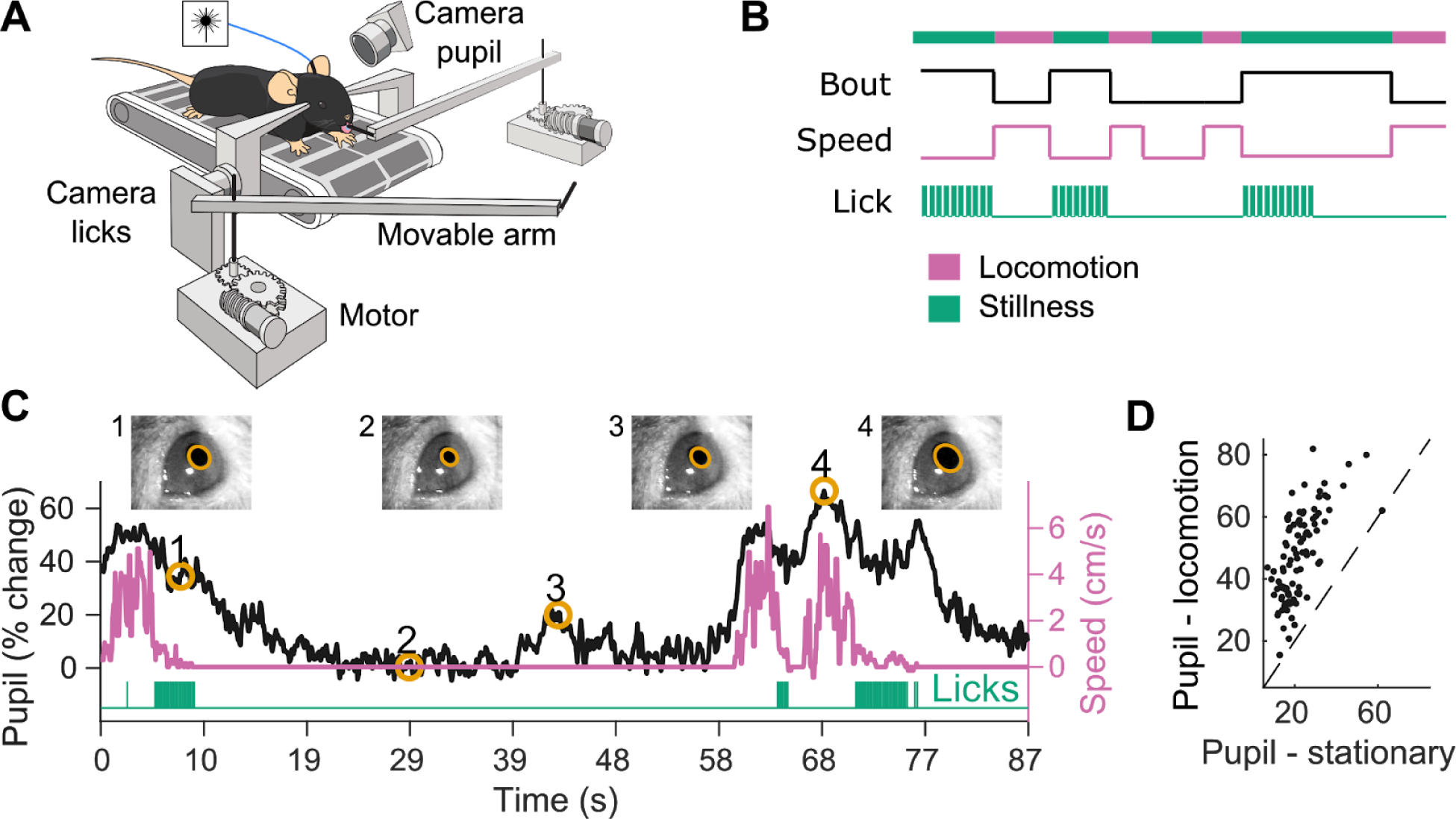
Tracking pupil-linked arousal during a foraging task for head-fixed mice. (A) A mouse placed on a treadmill exploited one of two resource sites materialized by movable arms. The mouse switched between sites by running a set distance on the treadmill during which time the site in front moved away and the distal one moved into place. Licks and the pupil were tracked by two different cameras and locomotion was monitored by the rotary encoder of the treadmill. (B) The task consisted of periods of locomotion and stillness. A behavioral bout was defined as the time spent at a given site. (C) Example pupil size (major axis in black), treadmill activity (pink), and lick time (green) from one experimental session. Pictures of the eye are shown at the time points indicated by the numbers on the pupil trace. (D) Average pupil size during periods of high locomotion (speed > 2.5 cm/s) versus stillness (speed < 0.5 cm/s). Each dot represents the average for each session (n = 96 sessions from 9 mice). See also Figure S1.

### Optogenetic activation of DRN 5-HT neurons increases pupil size

We trained mice that expressed Cre-recombinase under the control of the promoter of the 5-HT transporter (SERT-cre) and their wild-type littermates (WT). The DRN of both groups was infected with a viral vector containing Cre-dependent channelrhodopsin-2 and implanted with an optical fiber above the site of infection [28,29](Figure 2A). Post-hoc histological analyses confirmed ChR2-eYFP expression restricted to the DRN in SERT-Cre animals (Figure 2B & S2A,B; 141 ± 57 infected 5-HT neurons per animal) and no expression in WT controls (Figure S2C).

**Figure 2.**
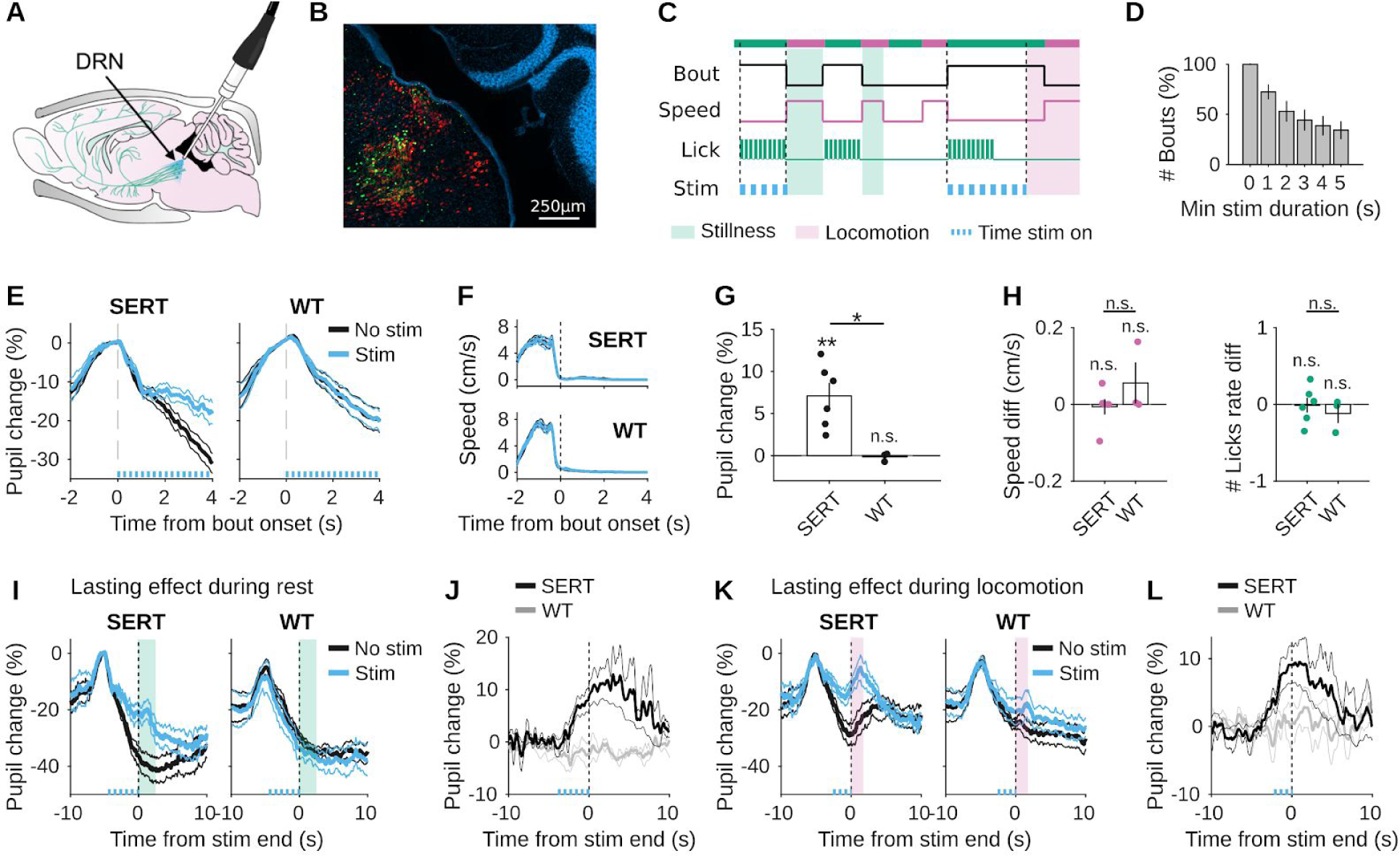
Optogenetic activation of DRN 5-HT neurons increases pupil size (A) Sagittal view of a mice brain illustrating the optogenetic approach. DRN neurons are infected with AAV2/9-Dio-ChR2-EYFP. In transgenic SERT-Cre mice (n = 6), 5-HT neurons express ChR2-YFP (green cells) and can be photoactivated with blue light (473 nm) delivered through an optical fiber implant. The same approach is used for WT littermates (n = 3), which do not express ChR2-YFP. (B) Fluorescence image of a parasagittal section showing 5-HT neurons labeled in red (rabbit anti-5HT) and ChR2-eYFP expression (green) localized to the DRN with DAPI in blue (See also Figure S2). (C) Schematic drawing of the stimulation protocol. The photostimulation (10 ms pulses, 25 s^-1^ at 5 mW for 30% of bouts) starts at the first lick and ends at running initiation or after 5 s of stillness. (D) Distribution of bout duration. For example, on average 72 ± 23% of bouts lasted for at least 1 s while 34 ± 27% of bouts lasted at least 5 s. (E) Time course of pupil responses aligned to the first lick of stimulated (blue) and non stimulated (black) bouts lasting at least 4 s for an example SERT-Cre and an example WT animal. (F) Average locomotor speed corresponding to the bouts in (E). (G) Average differences in pupil size between stimulated and non stimulated bouts for all mice were estimated during the last 0.5 s of stimulation. (H) Average difference in locomotor speed and lick rate between stimulated and non stimulated bouts for all mice. Speed was estimated in the same 0.5 s interval while lick rate is estimated in the full bout. (I) Time course of pupil responses aligned to the end of the photostimulation for stimulated (blue) and non stimulated (black) bouts lasting at least 7 s for the same animals as in (E). Here, the photostimulation lasted 5 s and mice remained still for at least 2 s after the photostimulation ended. (J) Summary across mice of the difference in pupil responses between stimulated and non stimulated bouts (i.e, the difference between the blue and black traces in I). (K) Time course of pupil responses aligned to the end of the photostimulation for stimulated (blue) and non stimulated (black) bouts of less than 5 s duration for the same animals as in (E, H). Here, the photostimulation lasted at least 3 s and mice terminated the stimulation with running initiation. (L) Summary across mice of the difference in pupil responses between stimulated and non stimulated bouts (i.e, the difference between the blue and black traces in K). All pupil responses in (E, I, K) are baseline-corrected by subtracting the median pupil in a 500 ms window before the first lick. Solid lines indicate the mean across sessions and thin lines the SEM. In (G,H) error bars indicate SEM. See also Figure S3.

We randomly selected 30% of behavioral bouts to stimulate DRN 5-HT neurons and compared bouts with and without stimulation over a total of 96 sessions (Figure 2C,D). We observed a significant increase in pupil size during photostimulation in SERT-Cre mice (mean ± SD, 7.1 ± 3.7%, p = 0.0058), but not in WT (−0.16 ± 0.50%, p = 0.64). This difference was also significant when comparing SERT-cre and WT mice (p = 0.0149). The effect of photosimulation could not be explained by a change in locomotor speed during the bout (SERT-Cre: p = 0.7698, WT: p = 0.4074, SERT-Cre vs WT: p = 0.2174), nor by a change in lick rate (SERT-cre: p = 0.9115, WT: p = 0.4448, SERT-cre vs WT: p = 0.5323; Figure 2E-H, Figure S3). These effects were sustained for a few seconds after the end of photostimulation, whether mice remained still (Figure 2 I,J, the effect lasts for 6.9 s, p < 0.01; see STAR Methods) or started to move (Figure 2K,L, the effect lasts for 5.1 s, p < 0.01). Together, these results show that phasic stimulation of DRN 5-HT neurons modulates pupil size in an acute and lasting fashion, and that the stimulation effects are not tightly coupled with locomotion.

### The effects of DRN 5-HT photostimulation are not specific to locomotor states

Changes in pupil-linked arousal are highly dependent on locomotion [2,26,27] (Figure 1). Thus, DRN photostimulation might modulate the relationship between locomotor speed and pupil size. This modulation could involve multiplicative or additive effects, or both (Figure 3A).

**Figure 3.**
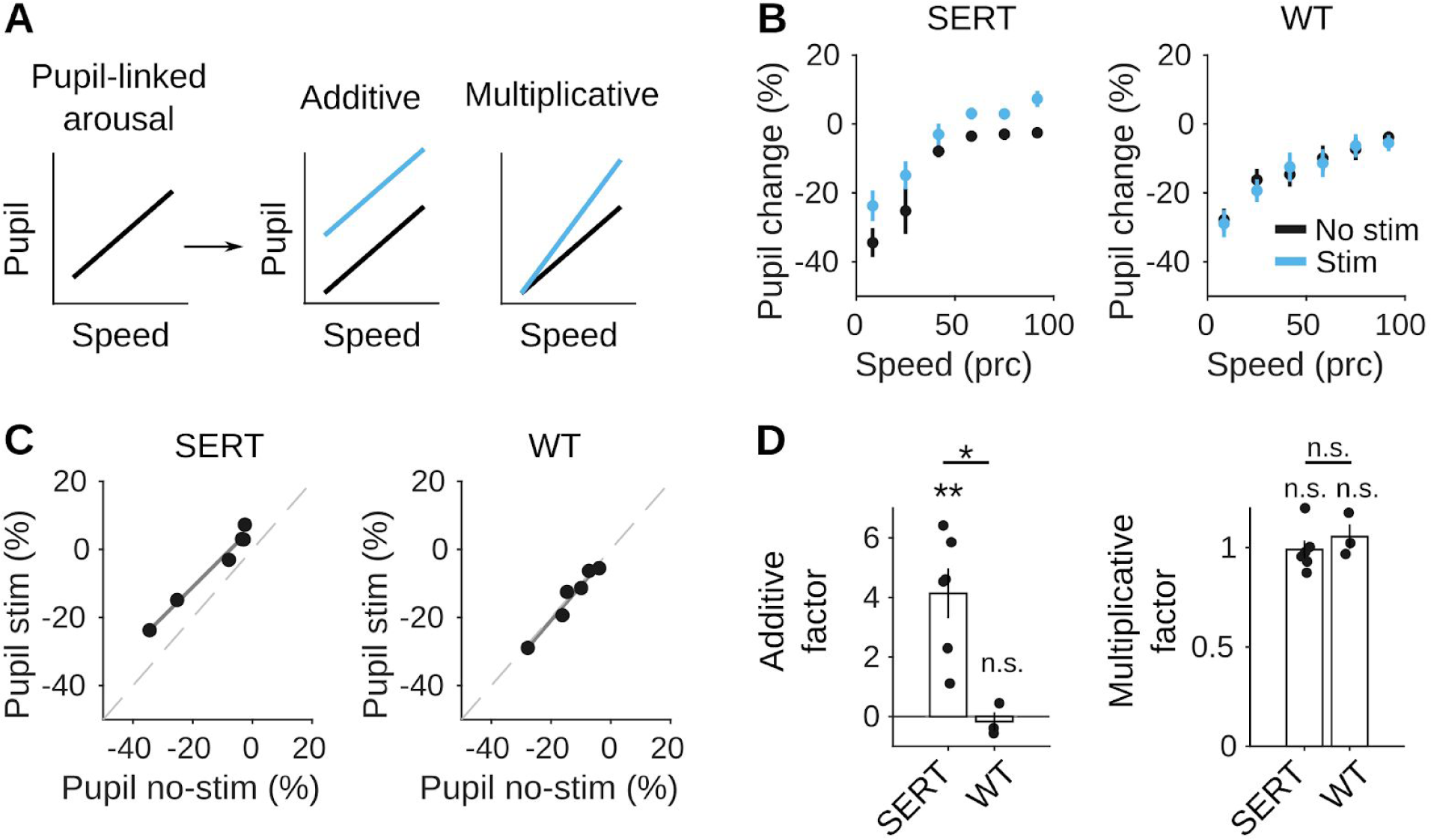
The effects of DRN 5-HT photostimulation are not specific to locomotor states. (A) Schematic drawing of hypothetical modulations of 5-HT neurons photostimulation on the relationship between speed and pupil size. (B) Pupil size as a function of speed averaged in a 4 s window after the end of the stimulation and hypothetical stimulation periods in non-stimulated bouts for a SERT-cre and WT example mice (mean ± SEM across sessions). (C) Mean pupil responses of the SERT-cre and WT mice in (B) after stimulated bouts (ordinate, corresponding to blue dots in (B)) vs. non-stimulated bouts (abscissa, black dots in (B)). Each dot is the mean pupil response corresponding to a given speed level. Grey lines are linear fits. (D) Average additive (i.e, intercept of linear fit in (C)) and multiplicative (i.e, slope of linear fit in (C)) factors for all SERT-cre and WT mice. Error bars indicate SEM.

To address this question, we built for each mouse an ‘input-output’ curve between speed and pupil response for stimulated and matched non-stimulated bouts (Figure 3B). Then, we characterized the additive and multiplicative modulations of photostimulation by performing linear regression between non-stimulated and stimulated bouts on the average pupil response to each level of speed (Figure 3C, see STAR Methods). The intercept of the fit indicates the additive shift in pupil response with photostimulation, whereas the slope of the linear fit describes how pupil size was scaled multiplicatively by the photostimulation. Hence, a positive intercept and a slope of one represents a purely additive effect, while a slope different from one and an intercept equal to zero represents a purely multiplicative effect.

We found that optogenetic activation of 5-HT neurons modulated the pupil size of SERT-Cre animals in an additive (p = 0.0043) but not a multiplicative manner (p = 0.8357; Figure 3D). The additive modulation was also greater in SERT-Cre than in wild-type mice (p = 0.0106), whereas the multiplicative factor was not significantly different between the two groups (p = 0.4287). This analysis shows that the increase in pupil-linked arousal by 5-HT neurons activation is not specific to the locomotor states of the animals.

### The effects of DRN 5-HT photostimulation depend on the level of uncertainty

Changes in pupil size have been linked to uncertainty, from noise in integration processes [30] to subjective uncertainty [4] and risk prediction errors or surprise [1,13,31,32]. In the foraging task, different levels of uncertainty can be achieved by varying the statistics of the environment (i.e., P_REW_ and P_DPL_) [25]. In the easy protocol where both P_REW_ and P_DPL_ are high, site depletions most often happen early in the bout and a few unrewarded licks are strong evidence in favor of site depletion. Hence, there is little uncertainty about whether or not the site is depleted. In the more uncertain protocol, lowering P_REW_ and P_DPL_ leads to many unrewarded licks, which are actually just unlucky attempts at a non-depleted site. Thus, by using protocol changes, we can manipulate the level of uncertainty.

To evaluate whether the increase in pupil size by 5-HT neurons activation scales with uncertainty, we examined the effect of photostimulation across different protocols (easy, medium and high uncertainty; one condition per session). We observed a graded effect of 5-HT activation on pupil size during the photostimulation (Figure 4A; 7.6 ± 5.4% p_easy_ < 10 ^-6^, 6.4 ± 4.0% p_medium_ < 10 ^-7^, 3.7 ± 5.8% p_hard_ = 0.0152, mean ± SD across sessions) and after the stimulation (Figure 4B; 10.3 ± 9.3%, p_easy_ < 10 ^-5^; 6.5 ± 5.3%, p_medium_ < 10 ^-5^; 4.4 ± 7.0%, p_hard_ = 0.0195). Specifically, pupils remained more dilated in protocols with low uncertainty than with high uncertainty, both during (p = 0.0260) and after the photostimulation (p = 0.0331). These results are consistent with previous works reporting that both behavioral effects of 5-HT transients [33] and changes in pupil sizes [4,13] depend on uncertainty. In particular, the decreasing effects of photostimulation with increasing protocol uncertainty could be consistent with the hypothesis that 5-HT transients signal a prediction error [24], a form of surprise reflected in the pupil response [13].

**Figure 4.**
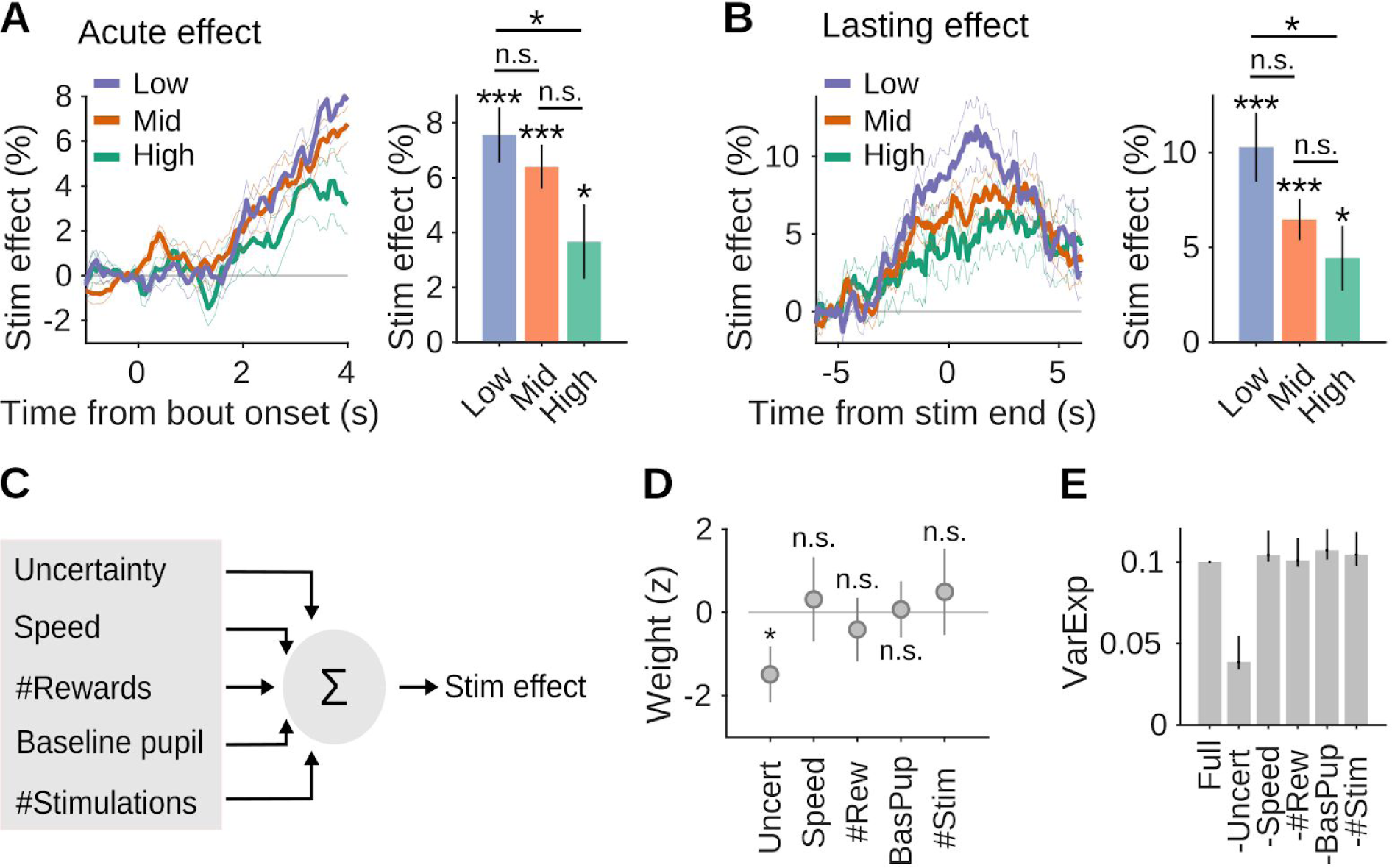
The effects of DRN 5-HT photostimulation depend on the level of uncertainty. (A) Average change in pupil responses (left) of SERT-cre mice between stimulated and non stimulated bouts lasting at least 4 s aligned to the first lick, and summary statistic (i.e, average difference in pupil size between stimulated and non stimulated bouts; right). Colors represent protocols with different levels of uncertainty (Low: P_REW_ = 90%, P_DPL_ = 30%; Medium, P_REW_ = 60%, P_DPL_ = 20%; High: P_REW_ = 45%, P_DPL_ = 15%). (B) Average change in pupil responses (left) of SERT-cre mice between stimulated and non stimulated bouts lasting at least 7 s and aligned to the end of the photostimulation and a summary statistic (right). (C) Schematics of the multivariate linear regression model. Speed is the average value in the session. Reward number indicates the mean number of rewards delivered during bouts. Baseline pupil is the median across bouts of the pupil size 500 ms before bout onset (the values used to baseline-correct the pupil when estimating the effects of the stimulation bout by bout). We also considered as a predictor the total number of stimulated bouts. (D) Weights obtained from fitting the data. Predictors were z-scored before fitting. Error bars indicate SEM. (E) Variance of the data explained by models where a given predictor is shuffled across sessions (−predictor name) compared with the model without any shuffling (Full).

In this paradigm, several alternative variables may interact with the effect of photostimulation on pupil size, such as the baseline pupil size, the average speed, the average number of rewards or the total number of stimulated behavioral bouts. We considered these alternative variables in a multivariate linear model to predict the effect of photostimulation on pupil size (Figure 4C, R^2^ = 0.1). The model confirmed that the uncertainty significantly affects the effect of photostimulation (negative coefficient significantly different from zero, Figure 4D, p = 0.03). To estimate the unique contribution of each of the predictors, we computed the explained variance using models where the predictor of interest was shuffled across sessions (Figure 4E). We found that the explained variance of the model only decreased (−6.0 ± 2.0 % compared to the full model) after shuffling the uncertainty predictor, confirming the importance of uncertainty in explaining the effects of the stimulation across sessions. Together, these results indicate that different levels of uncertainty predicted the magnitude of the 5-HT effect on pupil size, consistent with the idea that the effects of 5-HT transient on pupil-linked arousal are tightly linked to uncertainty.

Pupillary responses that coincide with transitions in brain states have been traditionally linked to the activity of the noradrenergic system from the locus coeruleus [6], and to a lesser extent to activity of the cholinergic system from the basal forebrain [34]. However, there has been no systematic investigation with other neuromodulators [17]. Here, by revealing a direct link between 5-HT neurons activation and phasic pupil dilation, our results expand our understanding on the neuromodulatory control of pupil-linked arousal. Our results also support the idea that noradrenaline and 5-HT may report closely related uncertainty-dependent signals [11,24]. Previous studies have shown reciprocal anatomical connections between these two major neuromodulatory systems with robust projections from the DRN to the locus coeruleus [35]. Yet, the nature of the interactions between 5-HT and noradrenaline remains to be understood, in particular whether they operate synergistically or play different functional roles in the context of unexpected uncertainty.

## ACKNOWLEDGEMENTS

We thank Anne Urai and Constanze Lenschow for helpful comments on the manuscript, and Michael Beckert for assistance with the illustrations. This work was supported by an EMBO long-term fellowship (F.C.; ALTF 461-2016), an AXA postdoctoral fellowship (F.C.), a Fundação para a Ciência e Tecnologia postdoctoral fellowship (D.R.; SFRH/BPD/119737/2016), a Marie-Curie postdoctoral fellowship (D.R.; H2020-MSCA-IF-2016 753819), the European Research Council Advanced Grant (Z.F.M.; 671251) and Champalimaud Foundation (A.R., Z.F.M.).

## AUTHOR CONTRIBUTIONS

F.C. and J.P.M. conducted the experiments, F.C. and D.R.designed and performed the analyses, F.C. and Z.F.M. designed the experiments, F.C and D.R wrote the paper, and J.P.M, A.R. and Z.F.M. reviewed and edited the paper.

## STAR METHODS

### Animal subjects

A total of 9 adult male and female SERT-cre mice (2-9 months old) were used in this study. All experimental procedures were approved and performed in accordance with the Champalimaud Centre for the Unknown Ethics Committee guidelines and by the Portuguese Veterinary General Board (Direco-Geral de Veterinria, approval 0421/000/000/2016). Mice were kept under a normal 12 hour light/dark cycle, and training occurred during the light period. Mice were water-restricted, and sucrose water (10%) was available to them only during the task. Mice were given 1 mL of water or 1 gram of hydrogel (Clear H2O) on days when no training happened or if they did not receive enough water during the task. Mice were housed individually after surgery and water-restriction started 7 to 10 days after surgery.

### Stereotaxic adeno-associated virus injection and cannula and head-plate implantation

Experimenters were blind to the mice’s genotype throughout the entire length of the experiment. Viral injection and fiber implantation were performed as described by Correia et al. [28]. Mice were anesthetized in an isoflurane induction chamber (2% for induction and 0.5–1% for maintenance with a 1.5% mixture with O2 and a flow rate of 0.8 L·min^-1^) and placed in the stereotaxic frame over a heating pad with the temperature set to 37°C. Animals’ eyes were covered and protected by the application of eye ointment (e.g., Vidisic, 2 mg/ml). For the injection, a craniotomy was performed over the cerebellum (−4.7 AP). A glass pipette was loaded with the viral solution (AAV2.9.EF1a.DIO.hChR2(H134R)-eYFP.WPRE.hGH, Addgene viral prep # 20298-AAV9) and lowered to the DRN (−4.7 AP, -2.9 DV) with a 32° angle. A volume of 1.2 mL of viral solution was injected using a Nanoject III (Drummond) at 40 nL·s^-1^. Fifteen minutes after injection, the pipette was removed and an optical fiber (200 μm core diameter, 0.48 NA, 4–5 mm long, Doric lenses) was slowly lowered through the craniotomy so that the tip of the fiber was placed 200μm above the target spot. Structural glue (Super-bond C&B kit) was used to fix the fiber to the skull. Carprofen solution (100 mL) was administered subcutaneously to provide analgesia. For the head plate implantation, four additional craniotomies were performed slightly anterior to the lambda stitch and four screws (Antrin miniature specialities, 000-120×1/16) were tightened inside each craniotomy. Super-Bond was used to fix a 22.3 mm metal head plate to the screws. After surgery, mice were removed from the stereotaxic frame and returned to their home cage where they were monitored for several hours. Animals were given a recovery period of at least a week before starting behavioral training.

### Histology and quantification of infected neurons

To assess viral expression and localization of ChR2-eYFP and optical fibre placement, we used postmortem histology at the end of the experiments. Mice were deeply anesthetized with pentobarbital (Eutasil, CEVA Sante Animale, Libourne, France) and perfused transcardially with 4% paraformaldehyde (P6148, Sigma-Aldrich). After perfusion, the brain was removed, and fixed for 24 hours in 4% PFA solution. Following fixation, the brain was transferred to phosphate buffer solution (PBS). Coronal or sagittal sections (40 µm) were cut with a vibratome (Leica VT 1000 S) and used for immunohistochemistry. Slices were washed in PBS and then blocked and permeabilized 0.3% Triton/10% FBS for 2 h. Slices were then incubated in blocking solution with primary antibody rabbit anti-5-HT (Immunostar #20080) with 0.1% sodium azide at 1/2000 dilution. Incubation occurred at room temperature in the dark, for 36 hours. Afterwards, slices were washed in PBS and incubated for 2 hours (in the dark, at room temperature) with secondary antibodies Alexa Fluor 594 (red, goat anti-rabbit,ThermoFisher #R37117) and Alexa Fluor 488 (green, goat, ThermoFisher #A11001) at 1:1000 dilution. Slices were mounted in Mowiol mounting medium and DAPI and finally sealed with nail polish. Scanning images DAPI, GFP and Alexa Fluor 592 were acquired with a slide scanner fluorescence microscope (Slide Scanner Axio Scan Z1, Zeiss, Oberkochen, Germany) equipped with a digital CCD camera (AxioCam MRm, Zeiss) with a 20× objective. Previous laboratory literature using the same Cre-dependent optogenetic approach and the same mouse line, reported that 94% of ChR2-eYFP-positive neurons were serotonergic [29]. Slide scans were analysed in QuPath [36] where distinct YFP-positive cell bodies were manually counted (Figure S2). To assess the region of infection, we mapped slices comparing the DAPI staining with the Allen Mouse Brain Atlas. This was achieved using a section aligner software QuickNII (RRID:SCR_016854) to anchor a reference DAPI section to the corresponding location in Allen Mouse Brain Atlas [37]. Locations of remaining sections were obtained by adding or removing 40 µm in sequential order from the reference section. The coordinates of all sections were individually estimated by matching to the high resolution Allen brain atlases (NeuN and NF-160 immunohistochemistry data; https://connectivity.brain-map.org/static/referencedata/).

### Experimental setup

Mice were head-fixed and placed on a linear treadmill with a 3D printed plastic base and a conveyor belt made of Lego small tread links. The running speed on the treadmill was monitored with a microcontroller (Arduino Mega 2560), which acquired the trace of an analog rotary encoder (MAE3 Absolute Magnetic Kit Encoder) embedded in the treadmill (speed measurements were computed as the analog signal from the rotary encoder smoothed by a median filter with a 250 ms time window and converted to cm/s). The treadmill could activate two movable arms in a closed-loop fashion via a coupling with two motors (Digital Servo motor Hitec HS-5625-MG). Water flowed through lick-ports glued at the extremities of each arm by gravity through water tubing. Water delivery was controlled by calibrated solenoid valves (Lee Company). Licks were detected in real time with a camera (Sony PlayStation 3 Eye Camera, 60 fps) located on the side on the treadmill using BONSAI [38]. The behavioral apparatus was controlled by microcontrollers (Arduino Mega 2560) and I/O boards (Champalimaud Hardware platform), which recorded the time of the licks, the running speed and controlled reward delivery and depletion according to the statistics of the task. Pupil videos were acquired at 15 fps using a monochromatic usb camera after the infrared filter was removed. Infrared lights were used to illuminate the pupil.

### Probabilistic foraging task

The task for head-fixed mice was adapted from a version developed for freely moving animals [25]. Here, mice collected water rewards by licking at a spout from either one of two ressource sites. At any given time, only one of the sites could deliver rewards, while the other was depleted. Reward delivery was probabilistic (given by P_RWD_) and each lick at a fresh site could trigger a stochastic site depletion (given by P_DPL_). When one site switched from fresh to depleted, the other necessarily switched from depleted to fresh. Thus, mice had to infer the state of the foraging sites to best decide when to switch between sites. Running on the treadmill activated the movement of the arms to allow mice to switch between sites. Mice performed under three different conditions of uncertainty: low uncertainty with P_REW_ 90% and P_DPL_ 30%; medium uncertainty with P_REW_60% and P_DPL_ 20%; and high uncertainty P_REW_ 45% and P_DPL_ 15%.

### Optogenetic stimulation

To optically stimulate ChR2-expressing 5-HT neurons, we used a laser emitting blue light at 473 nm (LRS-0473-PFF-00800-03, Laserglow Technologies, Toronto, Canada, or DHOM-M-473-200, UltraLasers, Inc., Newmarket, Canada). Light was emitted from the laser through an optical fiber patch-cord (200 μm, 0.22 NA, Doric lenses), connected to a second fiber patch-cord with a rotatory joint (FRJ 1×1, Doric lenses), which in turn was connected to the chronically implanted optic fiber cannula (M3 connector, Doric lenses). The power of the laser was calibrated before every session using an optical power meter kit (Digital Console with Slim Photodiode Sensor, PM100D, Thorlabs). During the foraging task, the optical stimulation (10 ms pulses, 25 s^-1^, 5 mW) was turned on during 30% of randomly interleaved bouts. Light delivery started after the first lick was detected, and lasted up to 5 s unless the animal started running, which interrupted the stimulation. Previous experiments validated that 5-HT neurons keep responding throughout the entire length of photostimulation [39] and that the photostimulation affected mice behavior [39–41].

### Pupil tracking

Pupil size and location were estimated using custom-made MATLAB (Mathworks, R2018a) scripts. We denoised each frame using a Wiener filter (using neighborhoods of size 5×5 pixels) and we applied lazy snapping [42] to segment each frame into background and foreground. The approach consisted of two steps. First, we separated the eye from the rest of the image. Then, we separated the pupil from the rest of the eye. Both procedures were performed based on pixel-seeds corresponding to different intensities (the eye is darker than the rest of the face and the pupil is the darkest object in the eye). Intensity thresholds for performing these operations were manually adjusted for each video/session by visually inspecting 20 random frames. Once the pupil was isolated, an ellipse was fitted. “Pupil size” was estimated as the major axis of the ellipse. To directly remove outlier estimations we took advantage of the slow time constant of pupil changes and applied a robust smoothing filter (linear, with a 250 ms window). To convert the pupil measurements (in pixels) to percent change relative to baseline, we normalized pupil size by the median 2% smallest values. These values mainly correspond to stationary periods. We assessed the quality of the tracking by visually examining 64 random frames of each video.

### Analysis of pupil response

To estimate the acute effects of the stimulation we compared, for each session, the median values of the pupil in non stimulated and stimulated bouts. Since pupil size greatly fluctuates throughout a session, we subtracted the median value of the pupil in the 500 ms before the first lick of each bout. The effects of the stimulation were then estimated as the difference between stimulated and non-stimulated trials in the 3.5-4 s after the beginning of the bout (for bouts lasting at least 4 s). We controlled for the robustness of the results on the acute effects of stimulation in different ways. First, we visually inspected whether the changes in pupil size were visible by eyes in the sessions (example in Figure S3.A) before any normalization or baseline subtraction. Second, we compared pupil size and locomotor speed in non-stimulated trials for SERT-cre and WT animals and found that they overlapped (Figure S3.B). We also checked the values of the baselines pupil for stimulated and non stimulated bouts for both SERT-cre and WT mice and found that baselines differences were not significantly different from zero (p = 0.58 for SERT-cre and p = 0.18 for WT animals, Figure S3.C). Then, we estimated directly the acute effects of the photostimulation without baseline subtraction and found results similar to Figure 2 (Figure S3.D). Finally, we also estimated the effects of the stimulation without baseline correction but z-scoring each session (Figure S3.E). We found again that the effect of the DRN stimulation on the pupil was consistent with the results reported in Figure 2 computed with baseline subtraction. Taken together all these results suggest that the validity of the reported effect does not depend directly on the fine details of the analysis. To estimate input/output curves between speed and pupil (Figure 3), we estimated the fluctuations in locomotor speed after the stimulation ends and the corresponding changes in pupil size (relative to baseline). In some bouts, mice started running quickly to reach the other site, while in some other bouts they slowly walked on the treadmill or stayed completely still for several seconds. We considered the speed and pupil in the 4 seconds following the end of stimulation for non-stimulated (the hypothetical end of the stimulation if it was applied) and stimulated trials. We computed the median pupil and the mean speed in this 4 s window for each bout. Then, to obtain an input/output curve for each session, we expressed the average speed in percentiles and calculated the corresponding median change in pupil size. Percentiles were used to allow comparison across sessions (mice may run more or less for some sessions). Input/output curves were averaged across sessions to estimate a mouse input/output curve for non-stimulated and stimulated bouts. Finally, we applied a linear fit for speed-matched pupil values to estimate the additive (intercept) and multiplicative (slope) effect of the stimulation for each mouse.

### Statistical tests

Results in the text are reported as means ± standard deviations across either mice or behavioral sessions. Behavioral sessions were considered to assess the correlation between locomotion and pupil size (Figure 1) and to quantify the effect of stimulation with varying uncertainty levels in the experimental protocol. We assessed statistical significance by performing t-tests (either to estimate whether the mean effect across mice was different from zero or to test the differences between SERT-Cre and WT mice). To estimate the acute effects of DRN stimulation in Figure 2G and 4A we considered the difference in pupil size between stimulated and non stimulated bouts between 3.5 s and 4 s after stimulus onset. To assess the duration of the lasting effects in Figure 2J,L, we shuffled stimulated and non-stimulated bouts within each session and we performed the same analysis as in Figures 2J-L. We then defined the duration of the effect as the point in time at which the value of the difference in pupil between stimulated and non stimulated bouts crosses the 99th percentiles of the shuffled version. To estimate the after stimulation effects in Figure 4B, we considered the difference in pupil size between non-stimulated and stimulated bouts in the 1 s following the end of the stimulation.

### Multivariate linear regression

To test the contribution of uncertainty and alternative variables on the effect of stimulation on pupil size, we performed a multivariate linear regression (using MATLAB “glmfit” function). The model predicted the magnitude of the stimulation effect session by session as a function of multiple predictors. Specifically, we predicted the percent difference between the median pupil for stimulated minus non stimulated bouts across 69 sessions. The analysis was restricted to bouts where the stimulation was at least 4 s. Our predictors were the baseline pupil (estimated as the mean across bouts of the median pupil in the 500 ms time interval before bout initiation), the total number of stimulated bouts, the level of uncertainty (values of 1, 2, 3 for the three different protocols used in the task, i.e., P_RWD_/P_DPL_= 90%/30%; 60%/20%, 45%/15%), the average speed in the session and the average number of rewards obtained in the session. As for the analysis in Figure 2, we checked that differences between the baselines for stimulated and non-stimulate bouts were not significantly different for all conditions (90%/30%, p = 0.72; 60%/20%, p = 0.87; 45%/15%, p = 0.96). Speed was also considered because of the tight relationship between pupil size and locomotion. Finally, the number of rewards was included as a predictor since 5-HT effects have been linked to reward valuation [33,43]. Predictors were z-scored before fitting the data. To estimate their unique contribution, we computed the explained variance of reduced models where each predictor was shuffled, one by one, across sessions and compared it with the explained variance of the full model. Results in Figure 4E represent the median of the explained variance and 25th and 75th percentiles values (100 shuffles).

## SUPPLEMENTAL INFORMATION

**Figure S1 related to Figure 1.**
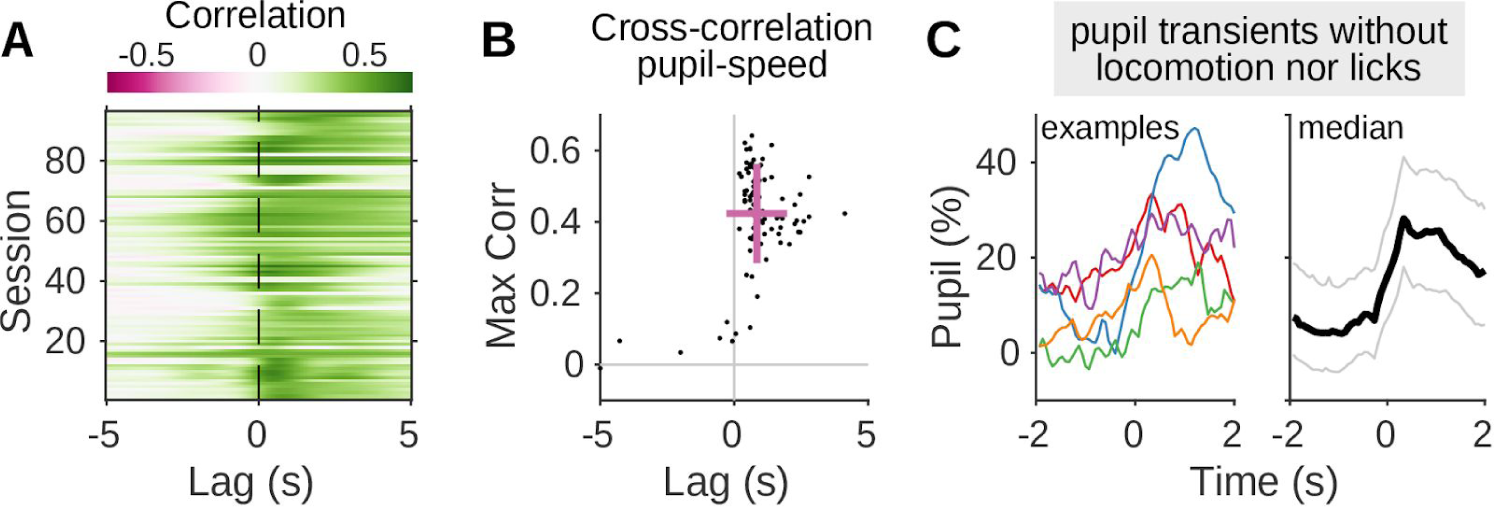
Pupil size fluctuations across locomotor states. (A) Session by session estimates of the cross-correlation between the pupil and locomotor speed. Color indicates the value of the correlation at that time lag. The increased correlations right after lag = 0 s indicate that pupil fluctuations sensitively follow changes in locomotor speed. (B) Summary statistics across sessions. Each point represents the maximum value of the cross-correlation (y-axis) and the corresponding time lag of the maximum (x-axis). Positive numbers indicate that the pupil is delayed compared to the speed. Violet lines represent mean±std across sessions. (C) Pupil transients appear even in the absence of locomotion and licks. These events were detected in periods of at least 15 seconds of null speed (0 cm·s^-1^), in the absence of licks and excluding 3 s at the beginning and end of these intervals (to exclude the slow drift at the end of a run or any pupil increase anticipating movement at the end of these intervals). Left panel: example traces. Right panel: median (black line) of these events and 25th and 75th percentiles (thin lines).

**Figure S2 related to Figure 2.**
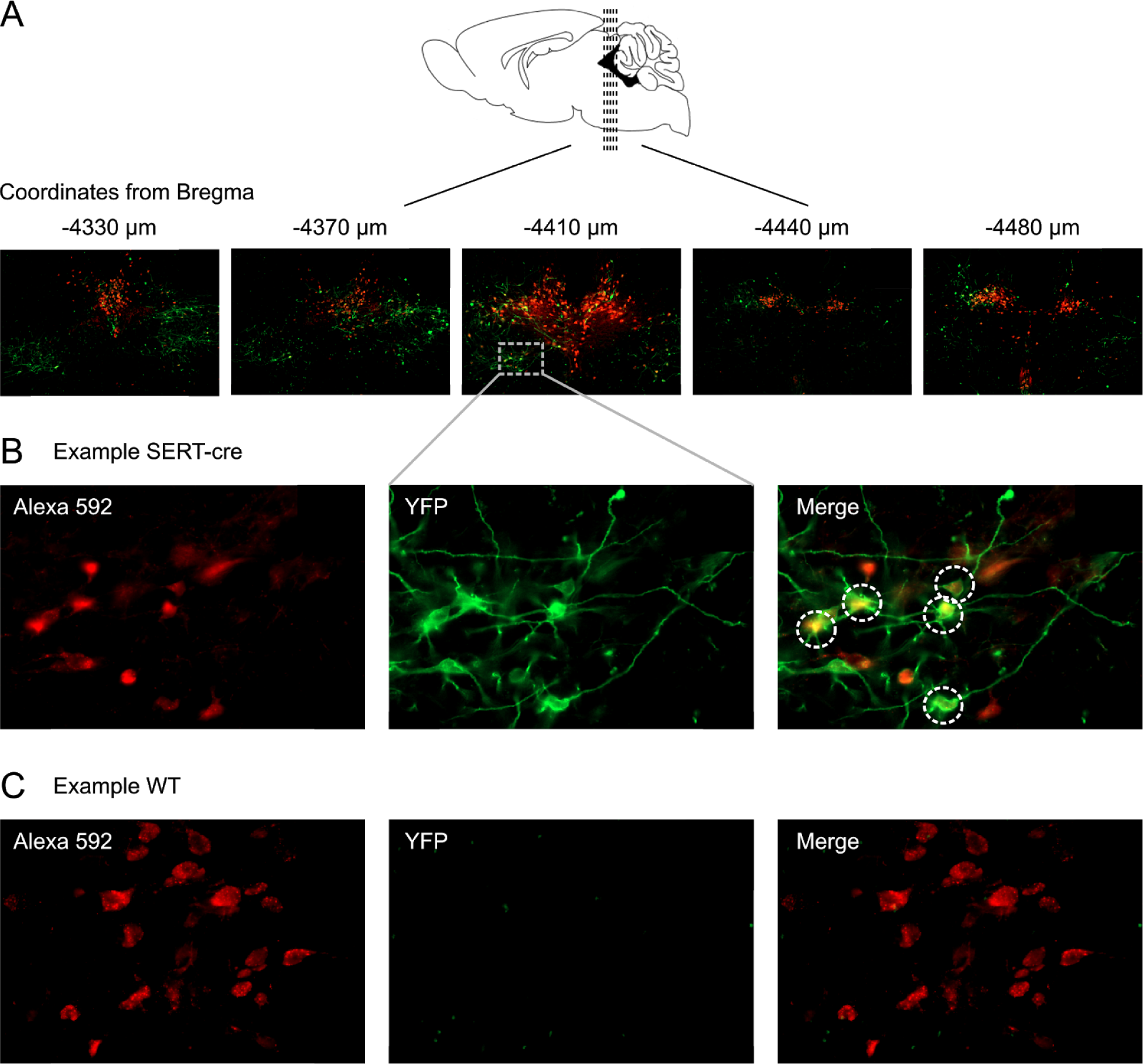
Assessment of viral expression and localization of ChR2-eYFP. (A) To evaluate viral expression and localization of ChR2-eYFP we acquired 20x z-stack images and examined and mapped coronal sections of the entire region around the DRN. (B) Images for Alexa Fluor 592 (left) and YFP (middle) were acquired to localize DRN 5-HT-positive neurons and fluorescent protein YFP respectively. We quantified the number of infected 5-HT neurons, identified by their expression of Alexa Fluor 592 (red) and YFP signal (green). The criteria used to consider an infected neuron were that both signals should be present in a cell body when background noise is null. The overlap of both signals will result in a distinct yellow color when merging the two images (right). Additional criteria were defined according to the physiological features of ChR2-eYFP and anti 5-HT. Since ChR2-eYFP is present in the cell membrane, clear green neurites should protrude from the candidate cell body while the 5-HT signal should be more concentrated in the soma. Using these additional criteria controlled for the presence of artifacts resulting from signal contamination between channels. (C) As a control for correct genotyping, the same method was applied to WT animals to verify that they did not express YFP and that in turn no cells were infected.

**Figure S3 related to Figure 2.**
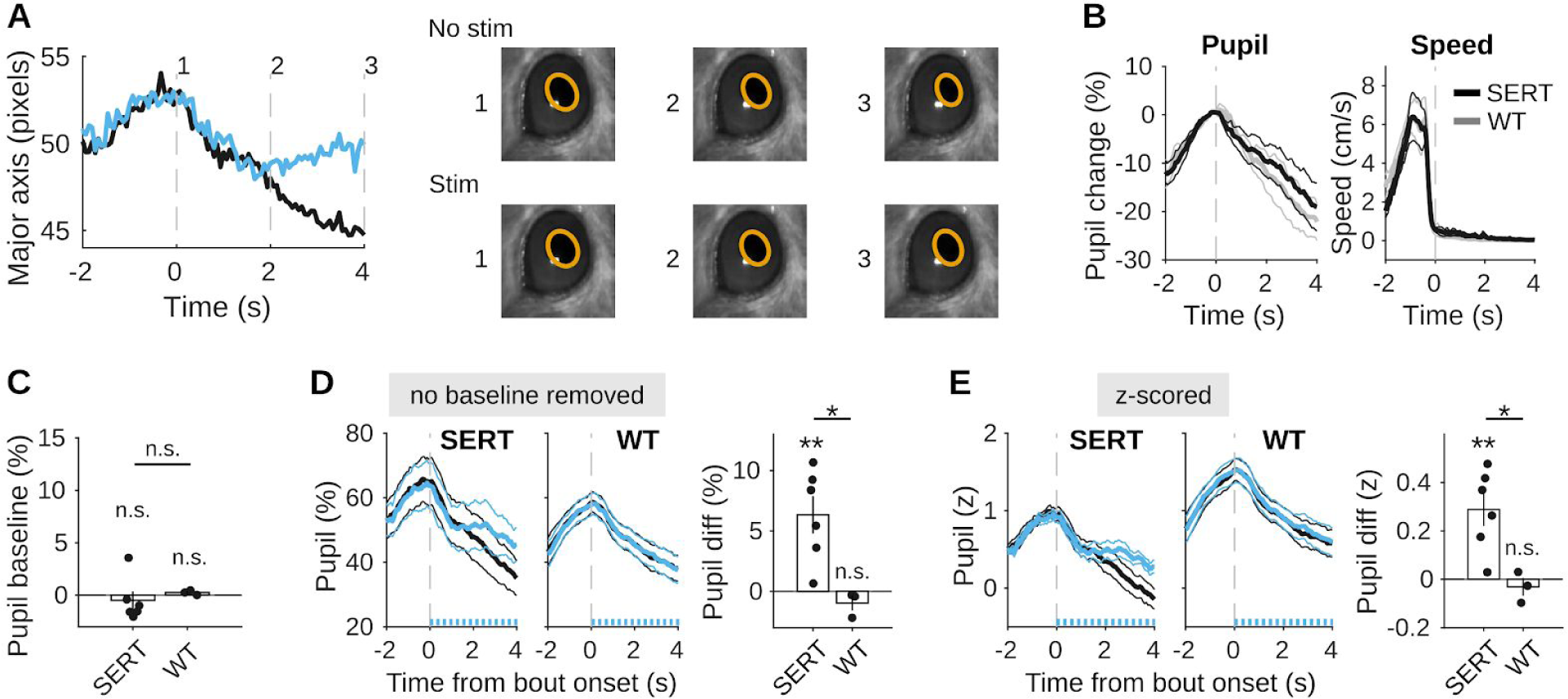
Robust effects of DRN 5-HT photostimulation on pupil size. (A) Left panel: median of raw pupil size values (pixels) in one example session aligned to bout onset. The blue line represents pupil size in stimulated bouts and black in non-stimulated ones. Right panel: images corresponding to the time points indicated by dash lines in the left panel. The difference in pupil size between stimulated and non stimulated bout in ‘3’ is visible “by eye”. (B) Average pupil and locomotor speed aligned to bout onset in non-stimulated bouts for both SERT-Cre and WT mice (mean ± SEM across animals). Traces overlap, indicating no major differences in baseline behavior between SERT-cre and WT mice. (C) Differences between the baseline values of the pupil for stimulated and non stimulated bouts for SERT-cre and WT mice. Differences are not statistically different from zero, suggesting that the effect of the photostimulation is not due to different baselines levels between stimulated and non stimulated bouts. (D) Left panel: Pupil traces aligned to bout onset for the same two example mice as in Figure 2. Differently from Figure 2, here the baselines are not subtracted, yet the differences in pupil size for stimulated and non stimulated bouts are still visible. Right panel: summary statistics of the acute effects of photostimulation without baseline subtraction. (E) Similarly to (D), here the pupil in each session was z-scored. Taken together, (D) and (E) show that the effects of the photostimulation do not depend on the specifics of the analysis.

